# A mutant bacteriophage evolved to infect resistant bacteria gained a broader host range

**DOI:** 10.1101/466383

**Authors:** Michal Habusha, Elhanan Tzipilevich, Sigal Ben-Yehuda

## Abstract

Bacteriophages (phages) are the most abundant entities in nature, yet little is known about their capacity to acquire new hosts and invade new niches. By exploiting the Gram positive soil bacterium *Bacillus subtilis* (*B. subtilis*) and its lytic phage SPO1 as a model, we followed the co-evolution of bacteria and phages. After infection, phage resistant bacteria were readily isolated. These bacteria were defective in production of glycosylated wall teichoic acid (TA) polymers, served as SPO1 receptor. Subsequently, a SPO1 mutant phage that could infect the resistant bacteria evolved. The emerging phage contained mutations in two genes, encoding the baseplate and fibers required for host attachment. Remarkably, the mutant phage gained the capacity to infect non-host *Bacillus* species that are not infected by the wild type phage. We provide evidence that the evolved phage lost its dependency on the species specific glycosylation pattern of TA polymers. Instead, the mutant phage gained the capacity to directly adhere to the TA backbone, conserved among different species, thereby crossing the species barrier.

## Introduction

Bacteriophages (phages) and their bacterial host display a constant evolutionary battle, leading to the emergence of mutations in both competing participants. Phage infection is initiated by the binding of phage tail proteins, to phage receptors located on the surface of the bacterial host (Rakhuba et al., 2010). The primary contact of the phage with the bacterial surface is often reversible, increasing the potential for the subsequent irreversible attachment to a second receptor, leading to DNA injection through the phage tail apparatus (Moldovan et al., 2007). The bacterial surface components show considerable variation among species, enabling highly specific interactions between phages and their target host (Koskella and Meaden, 2013). The Gram positive soil bacterium *Bacillus subtilis* (*B. subtilis*) contains surface polymers of teichoic acid (TA), a diverse family of cell surface glycopolymers containing phosphodiester-linked glycerol repeat units poly(Gro-P) (Sonenshein et al., 2002). The TA polymers can be either bound to the cytoplasmic membrane by a glycolipid anchor, lipoteichoic acid (LTA), or anchored to peptidoglycan (PG) through a N-acetylmannosaminyl β(1→4) N-acetylglucosamine linkage unit, wall teichoic acid (WTA). The WTA linkage unit is synthesized by a sequential action of TagO, TagA and TagB enzymes (Brown *et al.,* 2013). WTA is further divided into two forms: the major form, which comprises glycerol phosphate polymer, and the minor form, composed of a polymer of glucose (Glc) and N-acetylgalactosamine (GalNAc) (Brown et al., 2013; Sonenshein et al., 2002). The major TA form is decorated by Glc, and the Glc-WTA polymer is exploited by various *B. subtilis* phages as a binding receptor [e.g. (Yasbin et al., 1976; Young, 1967)].

Here we investigated the *B. subtilis* lytic phage SPO1, known to utilize the major Glc-WTA as its sole receptor (Yasbin et al., 1976). SPO1 is a large tailed bacteriophage from the *Myoviridae* family that harbors a genome of about 132 kb encased in a capsid of 87 nm in diameter (Parker et al., 1983; Stewart et al., 2009). *Myoviridae* phages exhibit a highly complex tail structure, consisting of a sheath, an internal tail tube, and a baseplate that holds the tail fibers (Fokine and Rossmann, 2014; Stewart et al., 2009). In general, the short tail fibers bind the Glc-WTA polymers, fastening the baseplate to the cell surface. Consequently, the baseplate opens up, the sheath contracts and the internal tube is pushed through the baseplate, penetrating the host envelope to inject the phage genome (Kostyuchenko et al., 2005; Leiman and Shneider, 2012).

Perhaps the simplest strategy employed by bacteria to acquire phage resistance is modifying phage receptor. If a phage is unable to interact with the host cell, infection will be subsequently prevented (Samson et al., 2013; Yasbin et al., 1976). To overcome this defense strategy, the phage, in turn, can modify the receptor binding protein (Samson et al., 2013). By studying an accelerated evolutionary process between *B. subtilis* and its lytic phage SPO1, we detected the frequent emergence of mutant bacteria resistant to the phage. In turn, we could isolate mutant phages capable of infecting resistant bacterial strains lacking the SPO1 receptor. Remarkably, these mutant phages also gained the capacity to infect non-host *Bacillus* strains, thus shedding light on how phages in nature can acquire new hosts.

## Results

### Isolation of *B. subtilis* mutants resistant to the SPO1 lytic phage

To study the co-evolution of *B. subtilis* and its phages, we aimed at isolating bacteria that became resistant to the SPO1 lytic phage. Following infection of *mCherry* labeled *B. subtilis* cells [MH1, derived from PY79 parental strain (Youngman et al., 1984)], a substantial number of cells survived, as revealed by the subsequent colony formation analysis (Fig. S1A). All the isolated colonies produced mCherry and were capable of growing over agar plates containing the SPO1 phage. However, variations were observed in the degree of phage resistance, with some of the colonies exhibiting full resistance, while others displaying partial phenotypes (Fig. S1B). Further examination of two fully (M1-M2) and three partially (M3-M5) resistant mutants showed their phenotype to be consistent when infection was monitored in liquid medium (Fig. 1A). Whole genome sequence (WGS) analysis revealed that the fully resistant mutants M1 and M2 contain mutations in *gtaC* and *gtaB*, respectively, whereas M3-M5 contain mutations in *tagE*, previously known as *gtaA* (Honeyman and Stewart, 1989) (Fig. 1B). M4 and M5 strains harbored the exact same mutation, hence only M4 was further investigated. Reintroducing the M4 mutation into the wild type strain resulted in recapitulation of the partial resistant phenotype (Fig. S1C). A mutant disrupted for the *tagE* (*ΩtagE*) gene was fully resistant to SPO1 (Fig. 1A), indicating that the obtained *tagE* mutations enabled partial functionality of the protein. Consistently, the level of glycosylation within the obtained mutants correlated with their level of resistance, as indicated by staining with fluorescently labeled lectin Concanavalin A (ConA) that interacts specifically with the glucosyl residues of the major WTA (Doyle and Birdsell, 1972) (Fig. 1D). The identified mutated genes are all part of the major WTA glycosylation pathway: GtaC and GtaB are required for synthesizing the sugar residue UDP-Glc that is then attached to the WTA polymers by TagE glycosyltransferase (Pooley et al., 1987; Sonenshein et al., 2002; Young, 1967) (Fig. 1C).

**Figure 1.**
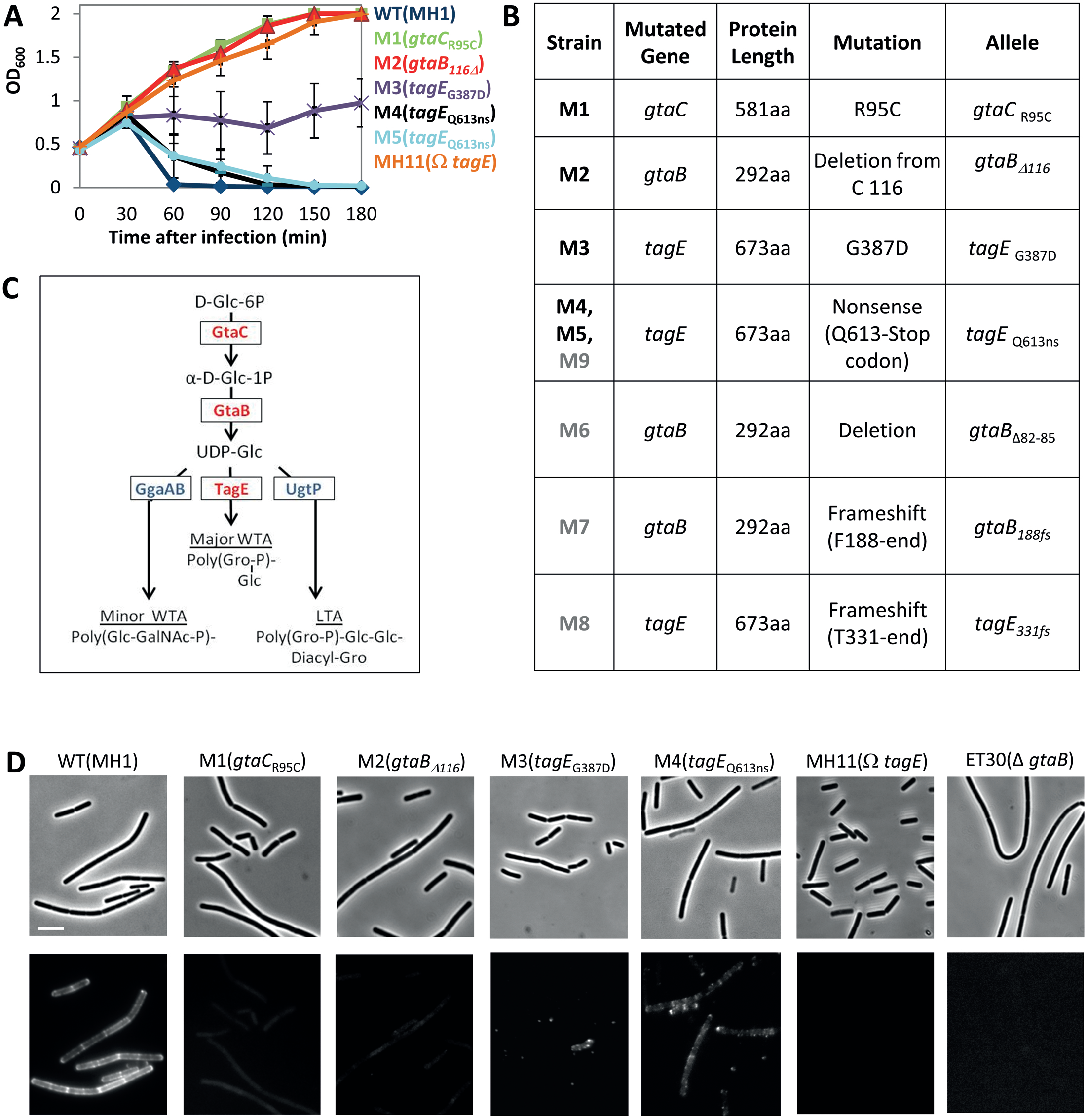
Phenotypes of phage resistance displayed by the isolated bacterial mutants. A. Lysis assay of WT (MH1) and the indicated mutant strains infected with SPO1 (MOI=10) (t=0). Cell lysis was followed by OD_600_ at the indicated time points. M4 and M5 infection kinetic exhibits only slight delay in liquid medium. Shown are average values and SD of at least 3 independent experiments.
B. Phage resistant mutants M1-M5 were characterized by WGS, whereas M6-M9 mutants (highlighted in grey) were characterized by sequence analysis of *gtaB*, *gtaC* and *tagE* loci. The indentified mutations in each strain are listed. Alleles are indicated by amino acid positions.
C. Diagram showing the flow of UDP-glucose (UDP-Glc) in the production of the three TA pathways and the key enzymes involved in their synthesis in *B. subtilis.* Adapted from (Schneewind and Missiakas, 2014).
D. ConA-AF488 labeling of Glc-WTA of the indicated bacterial strains. Shown are phase-contrast (upper panels) and the corresponding fluorescent ConA-AF488 (lower panels) images. Scale bar, 4 μm.

To extend our findings, four additional resistant strains (M6-M9) were subjected to sequence analysis of the *gtaB*, *gtaC* and *tagE* loci. M6 and M7 harbored deletions in *gtaB*, while the remaining strains (M8-M9) contained mutations in *tagE*, with M9 displaying a mutation identical to M4 and M5 (Fig. 1B). Thus, it seems that the most prevalent strategy for acquiring phage resistance is by excluding glycosylation of the cell surface major WTA polymers, which are utilized as a receptor by SPO1, as well as by other *B. subtilis* phages (Yasbin et al., 1976; Young, 1967).

### SPO1 utilizes the minor WTA for cell surface attachment

Since SPO1 receptor molecules were altered in the various resistant strains, we examined phage attachment to the surface of the partial and the fully resistant mutants. Phages were stained with the nucleic acid dye 4′,6-diamidino-2-phenylindole (DAPI), incubated with the different bacterial strains, and binding was examined by fluorescence microscopy. As expected, SPO1 was deficient in attachment to M1(*gtaC*) and M2(*gtaB*) (Fig. S2A). Nevertheless, SPO1 could efficiently adsorb to M3(*tagE_G387D_*) and M4(*tagE_Q613ns_*) *tagE* mutant strains, and even to MH11(*ΩtagE*) that lacks TagE (Fig. S2A). Similar results were obtained with classical attachment assay detecting free unadsorbed phages (Ellis and Delbruck, 1939) (Fig. S2B). Thus, SPO1 can attach to *B. subtilis* mutants lacking its known receptor, without causing their lysis (Fig. 1A), suggesting that an additional SPO1 binding component exists. To examine this possibility, we tested the capacity of SPO1 to bind and lyse mutants in the additional TA elements, LTA (*ΔugtP*) and minor WTA (*ΔggaAB*) (Jorasch et al., 1998; Lazarevic et al., 2005) (Fig. 1C). No effect was observed with the *ugtP* mutant, whereas a mutation in *ggaAB* largely reduced SPO1 binding (48%) (Fig. S2C). Accordingly, cell lysis kinetics of *ggaAB* mutant was slower and efficiency of plating (EOP) reduced (Fig. S2E and S2G). These results show that SPO1 attachment to the *ΩtagE* mutant is due to binding to the minor WTA. Although this attachment does not lead to a productive infection, it seems to facilitate SPO1 adsorption to the major form. Thus, similar to other phages, such as SPP1, K20 and T5 (Baptista et al., 2008; Heller and Braun, 1979; Silverman and Benson, 1987), SPO1 may have two modes of binding to the host, a reversible and an irreversible one.

### Isolation of phage mutants able to infect resistant bacteria

We next aimed to isolate phages that can infect cells fully resistant to SPO1. Streaking M2(*gtaB*) resistant strain on a plate containing SPO1 resulted in the formation of two plaques in two independent events. Phages were isolated from the plaques and their sequence was determined by WGS. The plaques yielded two different SPO1 phages (SPO1-m1 and SPO1-m2), carrying different missense mutations in the same genes: *gp16.2* and *gp18.1* (Fig. 2A), indicating that the combination of both is necessary for infecting resistant bacteria. The product of *gp16.2* has similarity to a baseplate protein of the T4 phage, whereas *gp18.1* encodes a protein with similarity to Phi29 host recognition tail appendages, corresponding to SPO1 tail fibers (Habann et al., 2014; Stewart et al., 2009) (Fig. 2B). Thus, both phages contain mutations within the same genes, encoding phage structural elements that are most likely involved in host cell recognition.

**Figure 2.**
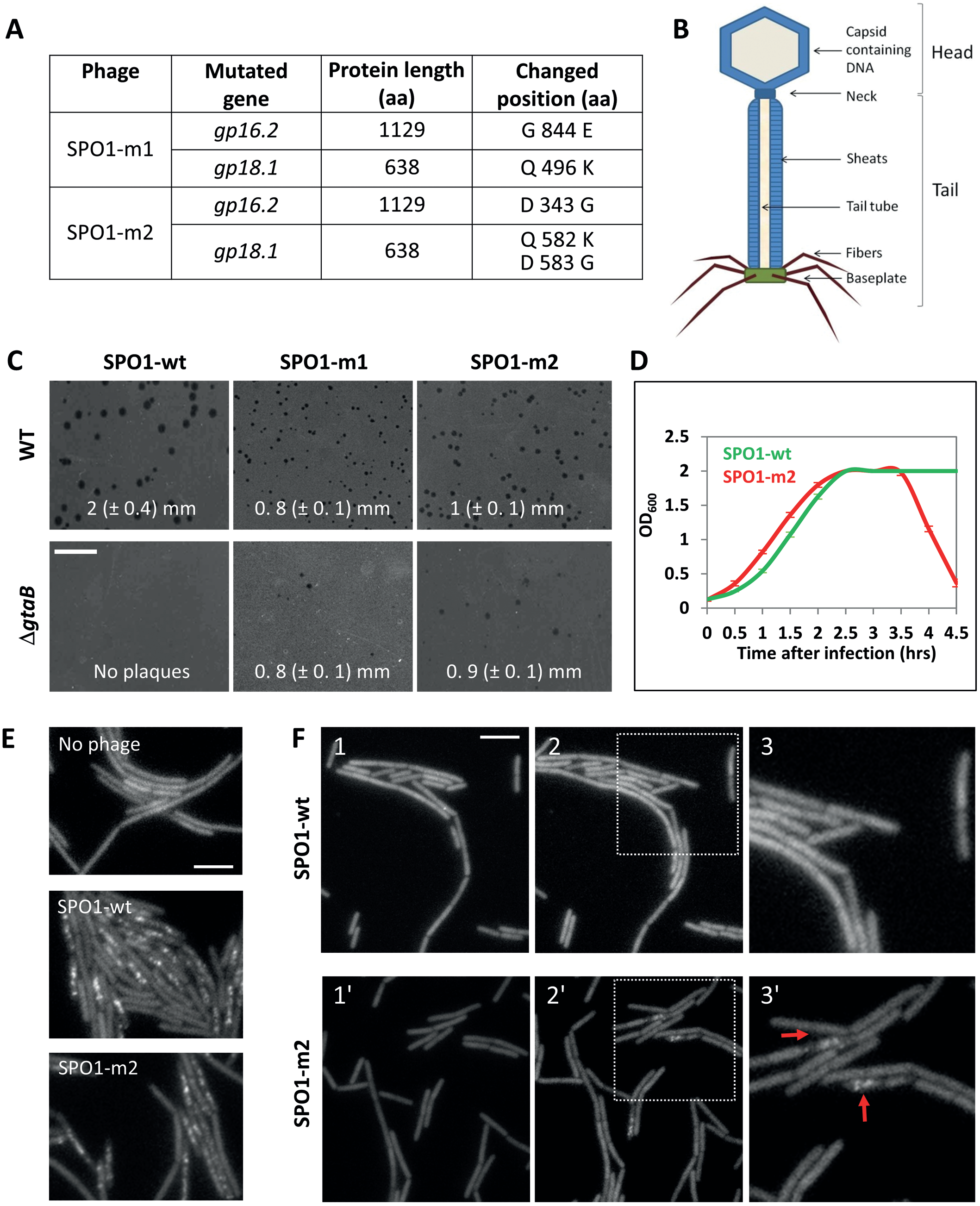
Isolation of SPO1 mutant phages that can infect resistant host cells. A. SPO1 mutant phages that could infect Δ*gtaB* (ET30) mutant cells were subjected to WGS. Mutation positions are listed.
B. Schematic representation of the general structural components of SPO1 phage. The phage tail consists of a contractile sheath, an interior tail tube through which the DNA is injected, and a baseplate. The baseplate carries the tail fibers, which serves as the phage host recognition receptor.
C. SPO1-wt, SPO1-m1 and SPO1-m2 phages were utilized to infect lawns of WT (PY79) and *ΔgtaB* (ET30) strains, and infection efficiency was detected by plaque formation using an agar overlay assay. The same concentration of phages was used for infection of both cell lawns (~10^4^ PFU/ml). Shown are representative plaque images and the average plaque diameter calculated from at least 30 plaques from 3 different independent experiments. Only unambiguous plaques from each experiment were considered for the analysis. Scale bar, 8 mm.
D. Δ*gtaB* (ET30) cells were infected with either SPO1-wt or SPO1-m2 (MOI=10) (t=0) and cell lysis was followed by OD_600_ at the indicated time points. Shown are average values and SD of at least 3 independent experiments.
E. Cells (ET8) harboring P_xyl_-*gfp*-*gp6.1* were grown in the presence of xylose (0.25%) and infected with SPO1-wt or SPO1-m2 (MOI=10). Shown are fluorescence images from GFP-GP6.1 expressing cells, uninfected (upper panel), or 1 hr post infection with SPO1-wt (mid panel) or SPO1-m2 (lower panel). Scale bar, 4 μm.
F. Cells (MH34) harboring *ΔgtaB* and carrying P_xyl_-*gfp*-*gp6.1* were grown in the presence of xylose (0.25%), infected with SPO1-wt or SPO1-m2 (MOI=10) (t=0), and followed by time lapse microscopy. Shown are fluorescence images from GFP-GP6.1 at t=1 hr (1, 1′), t=1.5 hrs (2, 2′) and a magnification of the framed regions in 2 and 2′ (3, 3′). Arrows highlight the formation of GFP-GP6.1 foci. Scale bar, 4 μm.

We next wished to characterize the infection capability of the mutant phages. SPO1-m1 and SPO1-m2 were used to infect the WT and the Δ*gtaB* strains (a complete deletion of *gtaB* was utilized in all following experiments). Both mutant phages were able to form plaques on WT cells, though plaque size was significantly smaller than those formed by SPO1-wt (Fig. 2C). Further, both mutant phages could form plaques on Δ*gtaB*, yet the number of plaques was reduced in comparison to plaques formed on a WT lawn (Fig. 2C). SPO1-m2 formed a higher number of plaques on Δ*gtaB* than SPO1-m1 (Fig. 2C), and therefore was solely utilized for further characterization.

To further demonstrate that SPO1-m2 can inject its DNA and propagate in cells lacking *gtaB*, we infected the cells with SPO1-m2 in liquid medium and could detect lysis of the population (Fig. 2D). Infecting additional TA mutants with SPO1-m2 revealed improved EOP on *ΔggaAB* cells (Fig. S2G), as well as a new capability to infect *ΩtagE* (compare Fig. S2E and Fig. S2F). To directly visualize SPO1-m2 entry into Δ*gtaB* cells, we constructed a strain harboring a reporter for SPO1 infection. SPO1 gene *gp6.1*, which encodes the phage capsid protein (Stewart et al., 2009), was fused to *gfp* (ET8) and introduced into the *B. subtilis* genome. Upon infection, GP6.1-GFP protein switches its localization from a diffused to a foci-like pattern, reporting phage invasion (Fig. 2E). Time lapse microscopy of Δ*gtaB* harboring the GP6.1-GFP reporter showed no appearance of foci upon infection with SPO1-wt. On the other hand, SPO1-m2 foci were clearly observed (Fig. 2F). These results substantiate the view that SPO1 mutant phages are able to infect Δ*gtaB* mutant cells.

### SPO1-m2 binding to resistant cells can be improved by experimental evolution

To gain insight into the strategy exploited by SPO1-m2 to infect Δ*gtaB* mutant, we investigated phage attachment to Δ*gtaB* cells, as well as to additional mutants in the TA pathway. The analysis indicated only a slightly improved binding of SPO1-m2 to Δ*gtaB* (compare Fig. S2C and Fig. S2D). The binding of SPO1-m2 to mutants in the additional TA pathways was similar to that of SPO1-wt (compare Fig. S2C and Fig. S2D). To better understand the infection strategy gained by the SPO1-m2 mutant phage, we attempted to improve its infectivity toward Δ*gtaB* cells by experimental evolution. Indeed, following 6 cycles of evolution, in which the phage was incubated with Δ*gtaB* host cells, a significant increase in SPO1-m2 adsorption and infectivity was observed (Fig. 3A-3B). WGS of SPO1-m2^∗^ isolate revealed the existence of two additional new mutations in *gp18.1* gene, and a missense mutation in *gp18.3* gene, encoding a putative tail base plate protein, in comparison to SPO1-m2 genome (Fig. 3C). Overall, our results suggest that the mutant phages acquired the capability to bind a surface component that cannot be recognized by SPO1-wt phage.

**Figure 3.**
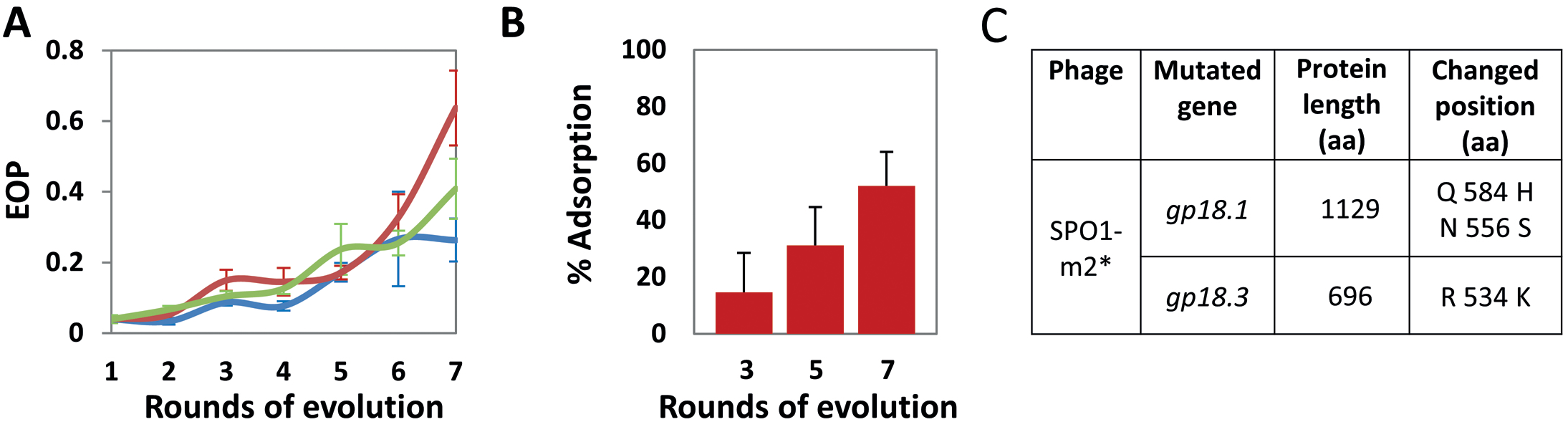
Evolution of SPO1-m2 yielded phages with increased infectivity of Δ*gtaB* cells. A. Three independent SPO1-m2 lysates were evolved on Δ*gtaB* (ET30) cells for 6 cycles, 1-7, whereby 1 is the starting population. EOF (efficiency of plating) on Δ*gtaB* (ET30) [number of plaques on ET30/ number of plaques on WT (PYT9)] was monitored for each cycle. The lysate with the best EOF (red line), termed SPO1-m2^∗^, was used for further characterization. Shown are average values and SD of at least 3 independent experiments.
B. Δ*gtaB* (ET30) cells were infected with SPO1-m2 from the indicated cycles of evolution in (H, red line) (MOI=0.1). Phage adsorption was monitored after 12 min by PFU/ml. Percentage phage adsorption was calculated as follows: (P_0_-P_1_)x100/P_0_, where P_0_ is the initial phage input in the lysate (PFU/ml), and P_1_ is the titer of free phages (PFU/ml) after cell infection. Shown are average values and SD of at least 3 independent experiments.
C. SPO1-m2^∗^ phage was subjected to WGS. The identified additional mutation positions are listed.

### *tagO* mutant cells are resistant to SPO1-m2

Interestingly, we could not detect the emergence of WT or Δ*gtaB* mutants resistant to SPO1-m2, neither from liquid culture nor from spotting high levels of phages on a lawn of cells (Fig. 4A-4B). We next searched for mutant bacteria, resistant to SPO1-m2 by transposon mutagenesis. Approximately 11,500 transposon mutated colonies of the Δ*gtaB* strain were streaked over plates containing SPO1-m2. The only two bacterial mutant colonies grown on the phage containing plates were mutated in the *znuC* gene, encoding a high affinity ABC zinc transporter, which imports zinc into the cell under conditions of zinc deprivation (Gaballa and Helmann, 1998). However, our analysis revealed that ZnuC does not function as a binding molecule for SPO1, and supplementation of excess Zinc suppressed the resistant phenotype (Fig. S3). Further, time lapse microscopy indicated that ZnuC acts to provide zinc to facilitate later stages of infection of both SPO1-wt and SPO1-m2 phages (Fig. S3E). Conducting additional genetic screens using chemical mutagenesis in an attempt to identify SPO1-m2 resistant mutants failed, and only mutants in *znuC* were repetitively isolated. This observation indicates that acquisition of such resistance is a rare event.

**Figure 4.**
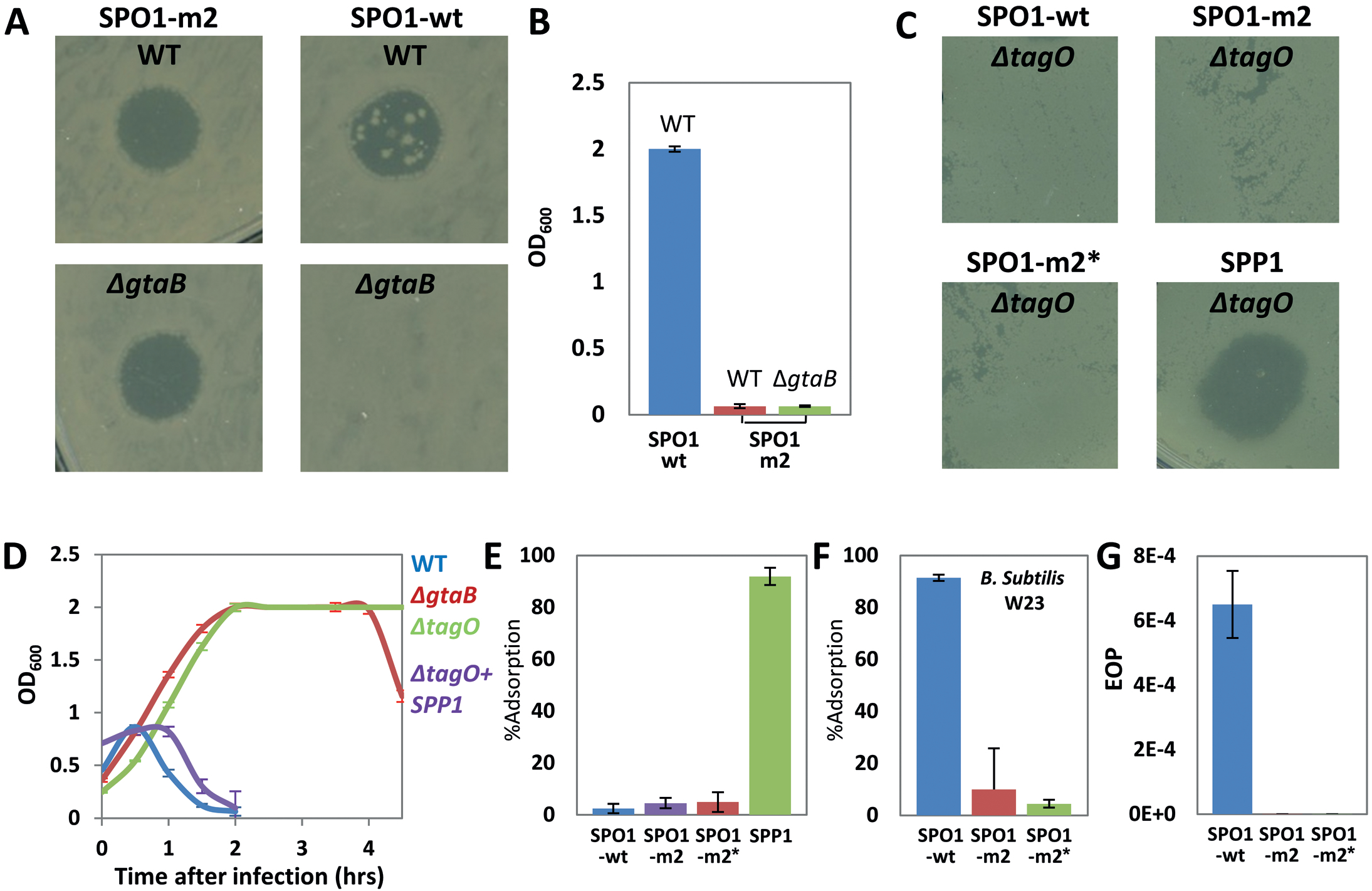
*tagO* mutant cells are resistant to SPO1-m2 phage. A. SPO1-wt and SPO1-m2 phages (10^8^ PFU) were spotted on lawns of *B. subtilis* (*B.s*) (PY79) or Δ*gtaB* (ET30) cells, and cell lysis was monitored after overnight. Spontaneous resistant colonies are seen within SPO1-wt lysis zone but not within SPO1-m2. Shown are representative images of at least 3 independent experiments.
B. WT (PY79) and Δ*gtaB* (ET30) cells were infected with either SPO1-wt or SPO1-m2 (MOI=10) (t=0), and OD_600_ was detected after 20 hrs of infection. Spontaneously arose resistant cells are capable of growing in the presence of the SPO1-wt phage. Shown are average values and SD of at least 3 independent experiments.
C. The indicated phages were spotted on lawns of Δ*tagO* (MH36) cells (10^8^ PFU). Cell lysis was monitored after overnight. Shown are representative images of at least 3 independent experiments.
D. Lysis assay of WT (PY79) Δ*gtaB* (ET30) and Δ*tagO* (MH36) strains infected with SPO1-m2 (MOI=10) (t=0). Cell lysis was followed by OD_600_ at the indicated time points. Δ*tagO* (MH36) strain, infected with SPP1 was followed for comparison. Shown are average values and SD of at least 3 independent experiments.
E. Δ*tagO* (MH36) cells were infected with the indicated phages, and phage (MOI=0.1) adsorption was monitored after 12 min by PFU/ml. Percentage phage adsorption was calculated as follows: (P_0_-P_1_)×100/P_0_, where P_0_ is the initial phage input in the lysate (PFU/ml), and P_1_ is the titer of free phages (PFU/ml) after cell infection. Shown are average values and SD of at least 3 independent experiments.
F. *B. subtilis* W23 cells were infected with the indicated phages (MOI=0.1), and phage adsorption was monitored after 12 min by PFU/ml. Percentage phage adsorption was calculated as follows: (P_0_-P_1_)×100/P_0_, where P_0_ is the initial phage input in the lysate (PFU/ml), and P_1_ is the titer of free phages (PFU/ml) after cell infection. Shown are average values and SD of at least 3 independent experiments.
G. *B. subtilis* PY79 and W23 cells were infected with the indicated phages at the same PFU and the number of plaques was monitored after 20 hrs. Shown is the number of plaques obtained on W23 cells divided by the number of plaques obtained on PY79 cells, i.e EOP (efficiency of plating) on W23 cells. Shown are average values and SD of at least 3 independent experiments.

Next, we combined several mutations in the TA assembly pathway in an attempt to decipher the SPO1-m2 binding molecule. Some reduction in SPO1-m2 infectivity was monitored, though it could not be eliminated (Fig. S4A-S4C). Nevertheless, deletion of the *tagO* gene, catalyzing the attachment of N-acetylglucosamine to undecaprenyl pyrophosphate, i.e the first step of the TA linkage unit synthesis (Brown *et al.*, 2013), conferred full resistance to SPO1-m2, as well as to SPO1-wt and SPO1-m2^∗^ (Fig. 4C-4E). Cells lacking *tagO* exhibit severe growth and morphology defects (Fig. S4D) (D’Elia et al., 2006), this could be the reason that mutants in *tagO* were not obtained following transposon mutagenesis. Nevertheless, *tagO* mutant cells were susceptible to SPP1 phage that utilizes a membrane protein (YueB) as the final receptor (Sao-Jose et al., 2004) (Fig. 4C-4E), signifying that the resistance to SPO1 is specifically due to the lack of TA.

In the absence of *tagO*, cells are lacking both the TA polymers and the linkage unit connecting them to the cell wall, placing each of these two components as a potential receptor for SPO1-m2. Since mutations in the assembly of TA polymers are lethal (Bhavsar et al., 2004; D’Elia et al., 2006), we sought to differentiate between these two possibilities by employing *B. subtilis* W23 strain, which utilizes the same TA linkage unit as PY79, but harbors TA polymers composed of a backbone of ribitol subunits poly(Rbo-P) instead of glycerol poly(Gro-P) (Anderson et al., 1977). W23 is sensitive to SPO1-wt, yet EOP is reduced in comparison to PY79 (Yasbin et al., 1976) (Fig. 4F-4G). Nevertheless, W23 was almost fully resistant to SPO1-m2 and SPO1-m2^∗^ (Fig. 4F-4G), indicating that TA polymers, rather than the common linkage unit, serve as a receptor for the mutant bacteriophages.

### SPO1 mutant phage gained a broader host range

While the glycosylation pattern of TA polymers largely varied among Gram positive bacteria, the TA polymer backbone is uniform among diverse species (Anderson et al., 1977; Illing, 2002). Therefore, we hypothesized that if SPO1-m2 interacts directly with the TA polymer backbone, it might exhibit a broader host range, capable of infecting non-host species, harboring poly(Gro-P)-based TA polymers (Brown *et al.*, 2013, *Illing *et al*, 2002*). To test this possibility, an array of *Bacillus* species were infected with SPO1-wt and SPO1-m2, and cell lysis was monitored (Fig. S5). Remarkably, indeed SPO1-m2 gained the capacity to lyse both *B. amyloliquefaciens* (*B. amylo*) and *B. pumilus*, close relatives of *B. subtilis*, which were not infected by SPO1-wt (Fig. 5A-5B and Fig. S5). Furthermore, a closer investigation of the *B. amylo* and *B. pumilus* interaction with SPO1-wt and SPO1-m2 by time lapse microcopy indicated the frequent lysis of both strains by SPO1-m2 but not by SPO1-wt (Fig. 5C and Fig. S6A), thus directly demonstrating the ability of the mutant phage to infect the new hosts.

**Figure 5.**
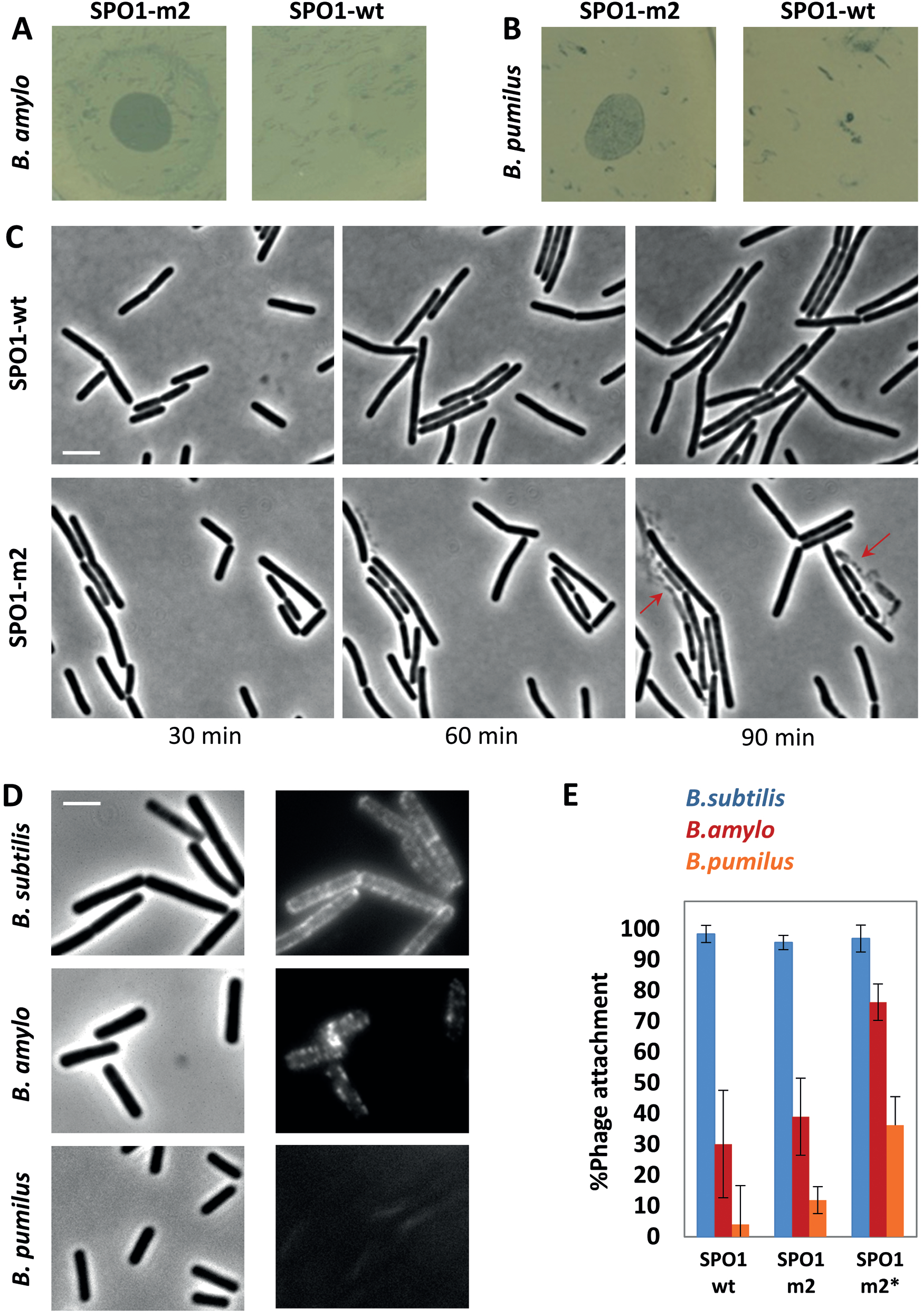
SPO1-m2 exhibits a broader host range. **(A-B)** SPO1-wt and SPO1-m2 phages were spotted on lawns of *B. amylo* (10A18) (A) and *B. pumilus* (B) cells (10^8^ PFU), and cell lysis was monitored after overnight. Shown are representative images of at least 3 independent experiments. **(C)** Infected *B. amylo* (10A18) cells were followed by time lapse microscopy. Shown are phase contrast images taken at the indicated time points post infection (t=0) with SPO1-wt (upper panels) or SPO1-m2 (lower panels) (MOI=10). **(D)** *B. subtilis* (PY79), *B. amylo* (10A18) and *B. pumilus* strains were stained with ConA-AF488. Shown are phase contrast (left panels) and AF488 fluorescence (right panels) images of the indicated strains. Scale bars, 1 μm. **(E)** *B. subtilis* (PY79), *B. amylo* (10A18) and *B. pumilus* strains were infected with the indicated phages (MOI=0.1), and phage adsorption was monitored after 12 min by PFU/ml on PY79 lawn. Percentage phage adsorption was calculated as follows: (P_0_-P_1_)×100/P_0_, where P_0_ is the initial phage input in the lysate (PFU/ml), and P_1_ is the titer of free phages (PFU/ml) after cell infection. Shown are average values and SD of 3 independent experiments. SPO1-m2 binding to *B. amylo* and *B. pumilus* was not significantly different from that of SPO1-wt (p=0.07 for *B. amylo*, and p=0.052 for *B. pumilus).* SPO1-m2 binding to both *B. amylo* and *B. pumilus* was significantly improved (p<0.01).

Comparing ConA staining of the newly invaded strains with that of *B. subtilis* (PY79) revealed a distinct pattern for *B. amylo* and the complete lack of signal from *B. pumilus* (Fig. 5D), signifying that both contain Glc-TA patterns different than that of *B. subtilis.* Monitoring phage binding to the new hosts by fluorescence microscopy and classical attachment assays denoted a slight increase in the attachment of SPO1-m2 in comparison to SPO1-wt that was significantly elevated by SPO1-m2^∗^ (Fig. 5E and Fig. S6B-S6C).

Taken together, our results show that the mutant phage gained the ability to infect resistant *B. subtilis* cells, and concomitantly acquired the advantage of invading new hosts.

## Discussion

The arms race between phages and their surrounding bacteria shapes the structure of bacterial communities in nature [e.g. (Samson et al., 2013; Suttle, 2007)]. Bacteriophages are typically specific to a single host species (Flores et al., 2011; Koskella and Meaden, 2013), and relatively little is known about their ability to acquire new hosts. Here we show that upon infection with a lytic phage, resistant bacterial mutants rapidly arose. In turn, mutant phages capable of infecting resistant bacteria evolved, and concomitantly gained the flexibility needed to cross the species barrier. The emergence of such adapted phages can provide a new conceptual understanding as of how phages in nature invade new hosts, and how the phage-host battle is accelerated.

Some phages of *Escherichia coli* (*E. coli*) were shown to gain the ability to infect resistant bacteria by acquiring mutations enabling the recognition of an alternative outer membrane receptor, homologous to the original one [e.g.(Drexler et al., 1989; Hashemolhosseini et al., 1994; Meyer et al., 2012)]. For example, mutations within the phage Ox2 allowed the recognition of the outer membrane protein OmpC, as an alternative to the original binding receptor OmpA (Drexler et al., 1989). Further, mutant phages that could recognize LPS molecules subsequently arose (Drexler et al., 1991). Here we characterized a SPO1 mutant phage that acquired the capability to infect bacteria resistant to the wild type phage, lacking its Glc-TA receptor. Based on our data, we propose that this mutant phage utilizes the TA polymer backbone as a receptor, bypassing the need for binding the glucosyl residues decorating these polymer chains. The capacity to recognize the TA polymers *per se*, which is less diverse than the glucosylation patterns (Illing *et al*, 2002, Lazarevic *et al.*, 2005), enables the mutant phage to bind and infect bacterial species harboring dissimilar glucosyltion patterns, but sharing a similar TA polymer backbone (Fig. 6).

**Figure 6.**
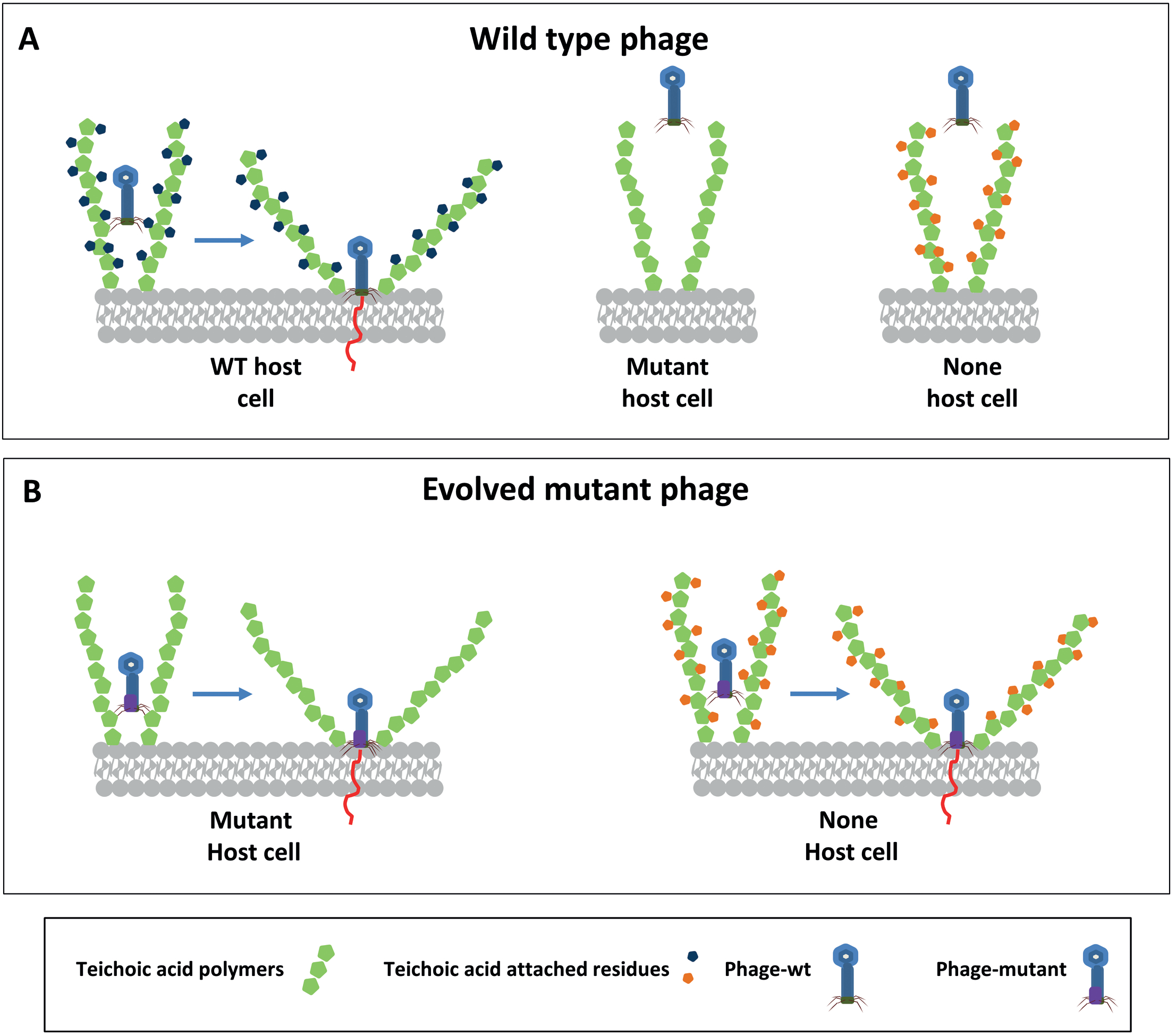
A model for host range expansion by phages. Based on our results, we propose the following model for host range expansion by phages infecting Gram positive bacteria: A. Wild type phage (i.e. SPO1-wt) recognizes the host species via binding to the Glc-TA polymers and subsequently injects its DNA into the cell. SPO1-wt phage cannot infect host cells lacking glucosyl residues or non-host species displaying a different TA glycosylation pattern.
B. The evolved mutant phage, capable of infecting resistant host (i.e. SPO1-m2), lost its dependency on TA glycosylation, and can directly adhere to the TA polymer backbone. The evolved phage can infect host cells having mutated or dissimilar TA glycosylation patterns.

In this study, we could easily isolate bacteria resistant to SPO1, with altered TA molecules. Since the majority of known *B. subtilis* phages utilize Glc-TA as a receptor for attachment and/or invasion into the host (Yasbin et al., 1976; Young, 1967), the emergence of “superresistant” bacteria seems an easy task. However, the preservation of Glc-TA in nature emphasizes the necessity of these surface molecules for bacterial physiology. TAs have been implicated in various cellular processes, such as structuring cell shape, cell division, colony formation and ion homeostasis (Brown et al., 2013; Mamou et al., 2017; Schneewind and Missiakas, 2014), yet the role of glycosylation of TAs is much less understood.

It has been recently shown that phage can broaden its host range to infect different bacterial species by evolving on an intermediate suboptimal host. This suboptimal host enables gradual adaptation toward a new host species. Such evolution was shown to occur in a multispecies bacterial community, like the gut microbiome, but could not be detected in a community composed of a pure species (De Sordi et al., 2017). Here we demonstrate that during co-evolution of a phage with its optimal-host species, an intermediate host evolved, which is resistance to the original phage, hence facilitating phage evolution toward the infection of new species.

We have previously provided evidence that phages can infect resistant bacteria that received the receptor from their sensitive neighboring cells (Tzipilevich et al., 2017). In this research, we describe a strategy by which phages gain mutations, enabling the infection of resistant bacteria in the absence of the phage natural receptor. As we showed, isolation of bacteria resistant to this mutated phage appears a difficult task, as *tagO* mutation causes severe growth defects, and defects in the synthesis of TA polymers has been shown to be lethal to the cells (Bhavsar et al., 2004; D’Elia et al., 2006). This observation raises the possibility that employing resistant bacteria to select for mutant phages can provide a promising strategy for a successful phage therapy.

## Experimental Procedures

### Strains and plasmids

All bacterial strains and phages utilized in this study are listed in Tables S1 and S2. Plasmid constructions were performed in *E. coli* (DH5α) using standard methods (Green et al., 2012) and are listed in Table S3. All primers used in this study are listed in Table S4.

### General growth conditions

Bacterial cultures were inoculated at 0.05 OD_600_ from an overnight culture grown at 23°C, and growth was carried out at 37°C in LB medium supplemented with 5 mM MgCl_2_ and 0.5 mM MnCl_2_ (MMB). Additional standard methods were carried out as previously described (Harwood and Cutting, 1990). For induction of the P_xyl_, xylose concentration was 0.25%.

### Phage lysate preparation

Phage lysate was prepared by adding approximately 10^9^ phages to 10 ml mid-log cells grown in MMB at 37°C, until the culture was completely cleared. Next, the lysate was filtered through a 0.22 μm Millipore filter. The number of phages was determined by PFU/ml. SPO1-m1 or SPO-m2 lysate was prepared from plaques with exponentially growing ET30 cells. The plate was incubated overnight at 37°C. Next, 7 ml of MMB was poured over the plate, incubated for an hour, and the liquid was collected and filtered through a 0.22 μm Millipore filter.

### Phage infection analysis

For liquid phage infection assays, the appropriate phage lysate (approximately 10^9^-10^10^ PFU/ml) was added to a mid-log (0.4-0.5 OD_600_) culture in 1:10 volume ratio (lysate: culture). Cells were grown in MMB at 37°C, and OD_600_ was measured every 30 min using spectrophotometer or microplate reader (Spark 10M, Tecan). Plaques formation analysis was carried out by adding 100 μl of phage lysate (approximately 10^3^-10^4^ PFU/ml), and 100 μl of bacterial culture (0.7 OD_600_) into 3 ml warm MMB containing 0.8% agar. The mixture was spread over MMB agar plate, incubated overnight at 37°C, and PFU/ml was determined. For preparation of phage containing plates, phage lysate (5 ml, approximately 10^10^ PFU/ml) was added to warm 250 ml MMB containing 1.6% agar, and the medium was poured into plates.

### Phage attachment assay

Indicated bacterial strains were grown in a liquid culture to OD_600_ 0.6, and phage adsorption to the cells was measured at 12 min post infection, by titrating the free phages present in the supernatant as previously described (Baptista et al., 2008). In brief, logarithmic cells were grown in MMB at 37 °C till 0.8 OD_600_, then 15 mM CaCl_2_ and 50 μg chloramphenicol/ml were added to the medium and cells were incubated for 10 min. Next, cells were infected with phages (10^7^ PFU/ml), and samples (0.5 ml) were collected at 12 min post infection, centrifuged for 1 min, and 50 μl of the supernatant was diluted, plated, and PFU/ml was determined.

### SPO1-m2^∗^ phage evolution

SPO1-m2 phage lysate (approximately 10^9^ PFU/ml) was added to a mid-log (0.5 OD_600_) culture of ET30 (Δ*gtaB*) in 1:10 (lysate: culture) volume ratio. Cultures were incubated overnight to ensure complete lysis. In the next day, the lysed culture was used to infect a fresh culture of ET30 (Δ*gtaB*). We estimate that each cycle included approximately 4 phage replication cycles.

### Phage spotting assay

Bacterial strains were grown in liquid culture to OD_600_ 0.8, 200 μl of cell were spread on LB plates in triplicates. The indicated phages (10^8^ PFU/ml, 10 μl) were spotted twice on each plate, and plates were incubated overnight at 37 °C

### Phage DNA labeling and microscope absorption assay

For DNA labeling, 100 μl of phage lysate (approximately 10^9^-10^10^ PFU/ml) were mixed with 2 μg/ml 4,6-Diamidino-2-phenylindole (DAPI) (Sigma) for 2 min. Labeled phages were dialyzed to remove excess dye using Slide-A-Lyzer MINI Dialysis Devices (Pierce Biotechnology). Lysate was placed in the dialysis tube and gently rolled inside 50 ml tube filled with MMB medium to remove the excess dye. Tubes were incubated over night (rolling shaker) in the dark at room temperature. For phage microscopic adsorption assay, 500 μl of mid-log growing cells were mixed with 15 μl labeled phages, incubated at room temperature for 10 min, and centrifuged for 2 min (8000 RPM) to pellet the cells. Supernatant was removed, and cells were suspended in the residual medium for further microscope analysis.

### Fluorescence microscopy

For fluorescence microscopy, bacterial cells (0.5 ml, OD_600_ 0.5) were centrifuged and suspended in 50 μl of MMB. For real time infection experiments, 1 ml logarithmic cells grown in MMB were infected with phages (100 μl of approximately 10^10^ PFU/ml). Culture was incubated at 37°C for 10 min, the infected cells were placed over 1% agarose pad and incubated in a temperature controlled chamber (Pecon-Zeiss) at 37°C. Samples were photographed using Axio Observer Z1 (Zeiss), equipped with CoolSnap HQII camera (Photometrics, Roper Scientific). System control and image processing were performed using MetaMorph 7.4 software (Molecular Devices).

### ConA labeling

Logarithmic cells grown in LB were suspended in 1 ml PBSx1 and washed once. Concanavalin A (ConA)-conjugated Alexa-Fluor 488 (ConA-AF488; Invitrogen) was dissolved in 0.1 M sodium bicarbonate (pH 8.3) to 5 mg/ml stock solution. ConA-AF488 was added to the cell suspension (200 μg/ml), and the mixture was incubated in the dark at room temperature for 30 min. Cells were washed 3 times, resuspended in PBSx1, and subjected to microscopy analysis.

### Isolation of phage resistant mutants

For spontaneous isolation of phage resistant mutants, 1 ml of mid-log (0.4-0.5 OD_600_) *B. subtilis* culture grown in MMB medium was infected with 100 μl of phage lysate (approximately 10^9^-10^10^ PFU/ml). Next, the infected culture was incubated at 37°C until clearance, and the lysate was spread over MMB plates without dilution. Plates were grown overnight at 37°C, and colonies were streaked over plates containing SPO1-wt. For isolation of phage resistant mutants, transposon mutagenesis of *B. subtilis* cells (MH30) was carried out as previously described (Pozsgai et al., 2012). Mutated colonies were grown on chloramphenicol containing plates, picked and streaked over plates containing SPO1-m2, and phage resistant colonies were isolated. Phage containing plates contained SPO1 lysate (5 ml, approximately 10^10^ PFU/ml) added to warm 250 ml MMB containing 1.6% agar.

### WGS analysis

Genomic DNA was extracted from mutant and parental strains using wizard genomic DNA purification kit (Promega). Libraries were prepared using Nextra XT kit (Illumina), and sequenced in a MiSeq sequencer (Illumina). The sequencing was pair end of 250bp X2. Quality assessment was carried out with the software FastQC (version 0.10.1). Sequence reads were aligned using NCBI *B. subtilis* PY79 genome (GenBank: CP004405.1). The sequence of the mutants was aligned with that of the parental strain. For phage genomic analysis, DNA was extracted using Phage DNA isolation Kit (Norgen biotek). Sequence reads of SPO1-wt were aligned using NCBI SPO1 phage genome (GenBank: Bacillus phage SPO1.NC_011421.1), and the mutant phages were aligned with the wild type phage.

## Acknowledgments

We thank David Rudner (Harvard University, USA), Dan Kearns (Indiana University, USA), Rotem Sorek (Weitzman institute Israel), Eric Brown (Mcmaster University, Canada) and Carlos São-José (Lisbon University, Portugal) for providing *B. subtilis* strains. We are grateful to Alex Rouvinski (Hebrew University, Israel) and members of the Ben-Yehuda laboratory for valuable comments.

## Author contributions

MH and ET have carried out the described experiments. MH, ET and SBY contributed to the conception and design of the study, interpretation of the data, and writing of the manuscript.

## Supporting Figure legends

**Figure S1.**
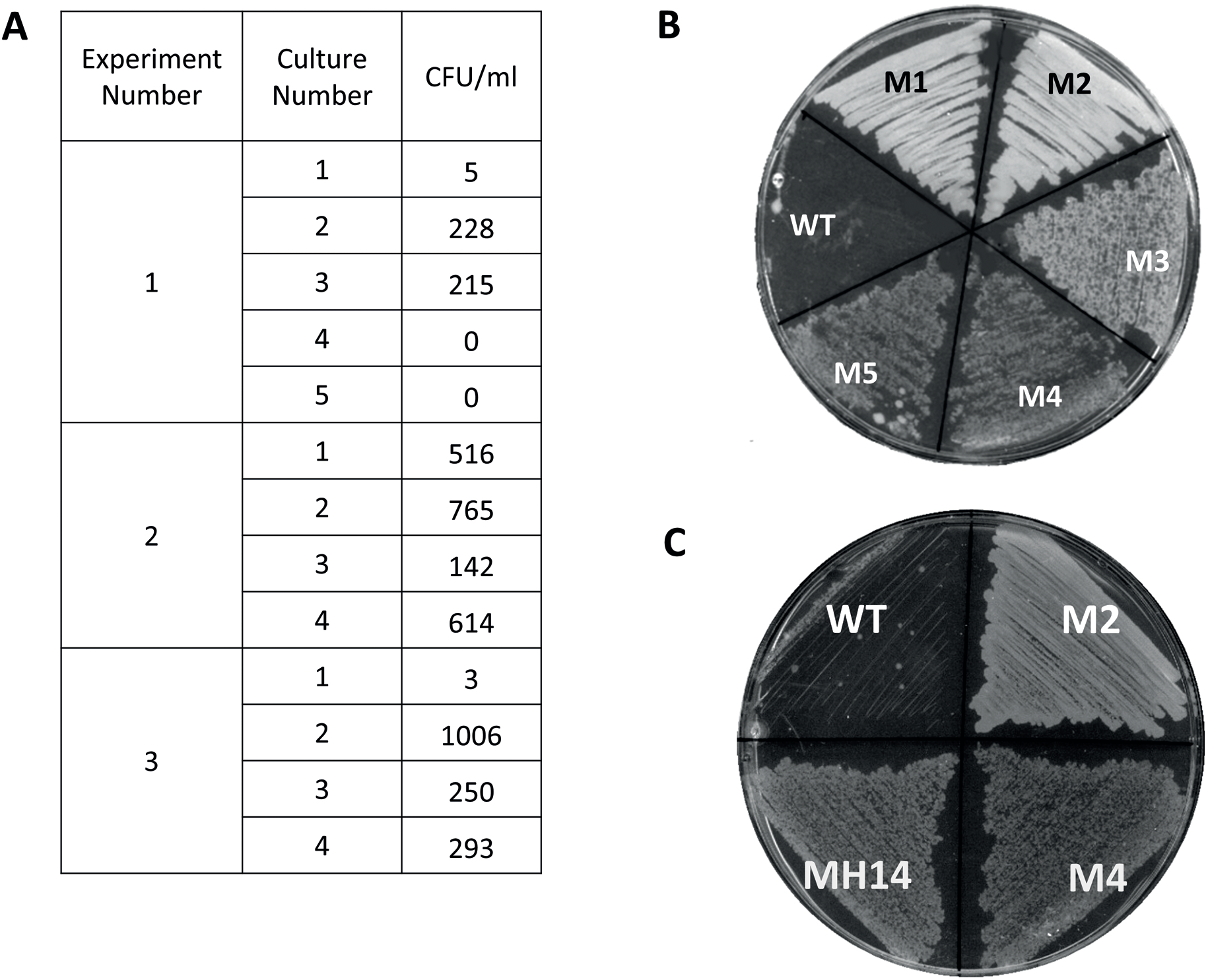
Isolation of bacterial strains resistant to SPO1. A. WT (MH1) strain was infected with SPO1 (MOI=10) and resistant colonies were scored. Shown are the numbers of phage surviving colonies in 3 representative biological repeats (out of 7 independent experiments). Each experiment included several parallel cultures, as indicated by culture number. Of note, the number of resistant colonies varied considerably between and within experiments.
B. WT (MH1) strain was infected with SPO1 and resistant colonies were isolated. Shown are various resistant bacterial colonies, M1-M5, streaked over an agar plate containing SPO1 phages. M1-M2 display full resistance, M3-M5 display partial resistance, as indicated by the formation of small plaques, and the WT (MH1) strain shows full sensitivity. The plate was photographed after 20 hrs of incubation at 37°C.
C. Strains WT(MH1), M2(*gtaB_116Δ_*), M4(*tagE*Q613ns) and MH14 (*tagE*Q613ns; a newly constructed M4) were streaked over an agar plate containing SPO1 phages. The plate was photographed after 20 hrs of incubation at 37°C.

**Figure S2.**
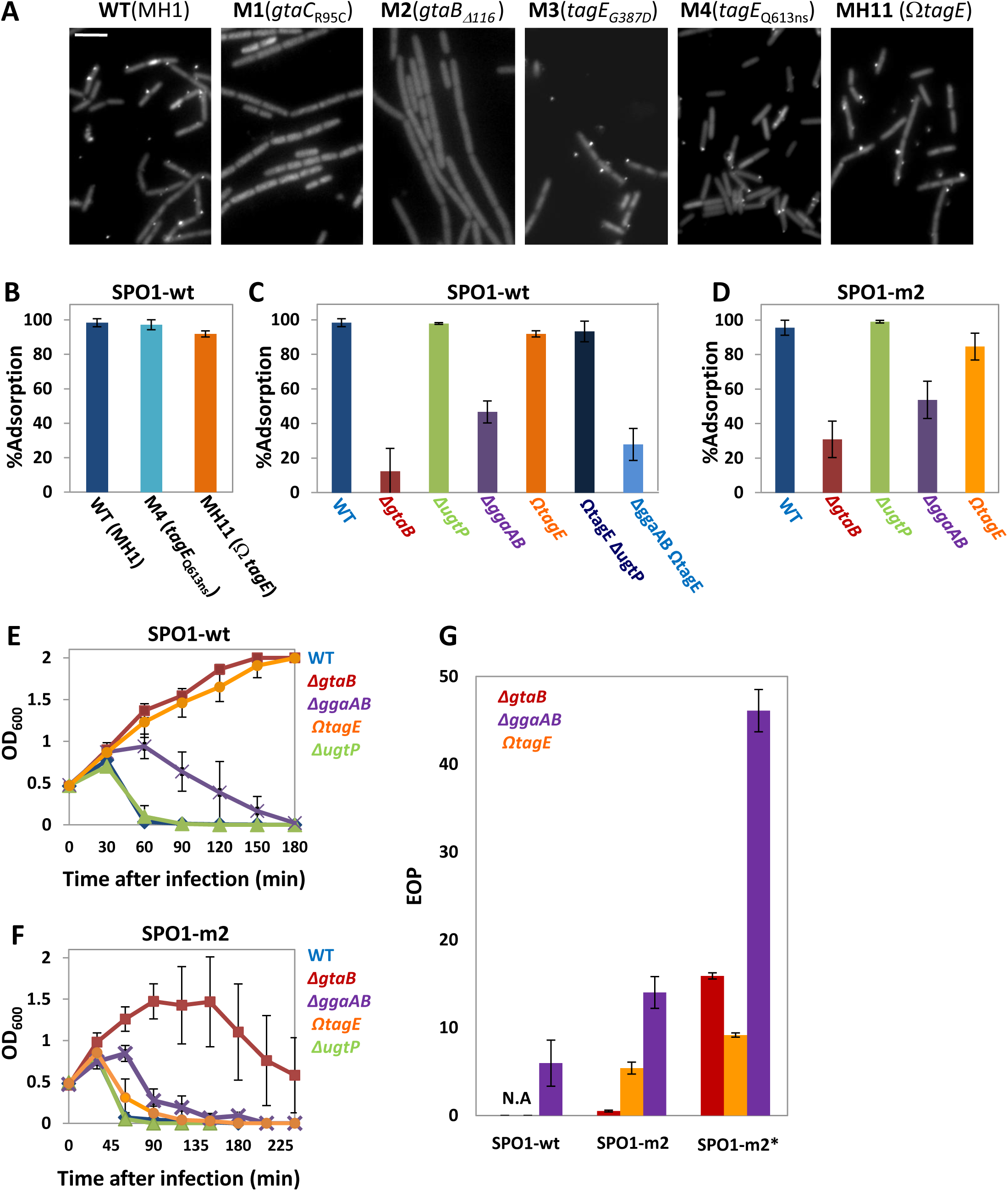
Phage attachment and infection of TA mutant bacterial strains. **(A)** SPO1 phages were labeled with DAPI and phage attachment to the indicated bacterial strains was observed by fluorescence microscopy. Scale bar, 4 μm. **(B-D)** The indicated bacterial strains were infected with SPO1-wt (B, C) or SPO1-m2 (D) (MOI=0.1), and phage adsorption was monitored after 12 min by PFU/ml. Percentage phage adsorption was calculated as follows: (P_0_-P_1_)×100/P_0_, where P_0_ is the initial phage input in the lysate (PFU/ml), and P_1_ is the titer of free phages (PFU/ml) after cell infection. Shown are average values and SD of at least 3 independent experiments. SPO1-m2 binds Δ*gtaB* significantly better than SPO1-wt (p<0.05). **(E-F)** The indicated bacterial strains were infected with SPO1-wt (E) or SPO1-m2 (F) (MOI=10) (t=0) and cell lysis was followed every 30 minutes by OD_600_. Shown are average values and SD of at least 3 independent experiments. **(G)** WT (PY79) or the indicated mutant strains were infected with the listed phages at the same PFU, and plaque number was monitored after 20 hrs. Shown is EOP (efficiency of plating) on the various mutants: [number of plaques on each mutant/number of plaques on WT]. Results are average values and SD of at least 3 independent experiments. N.A.-not applicable, no plaques were obtained on these bacterial mutants.

**Figure S3.**
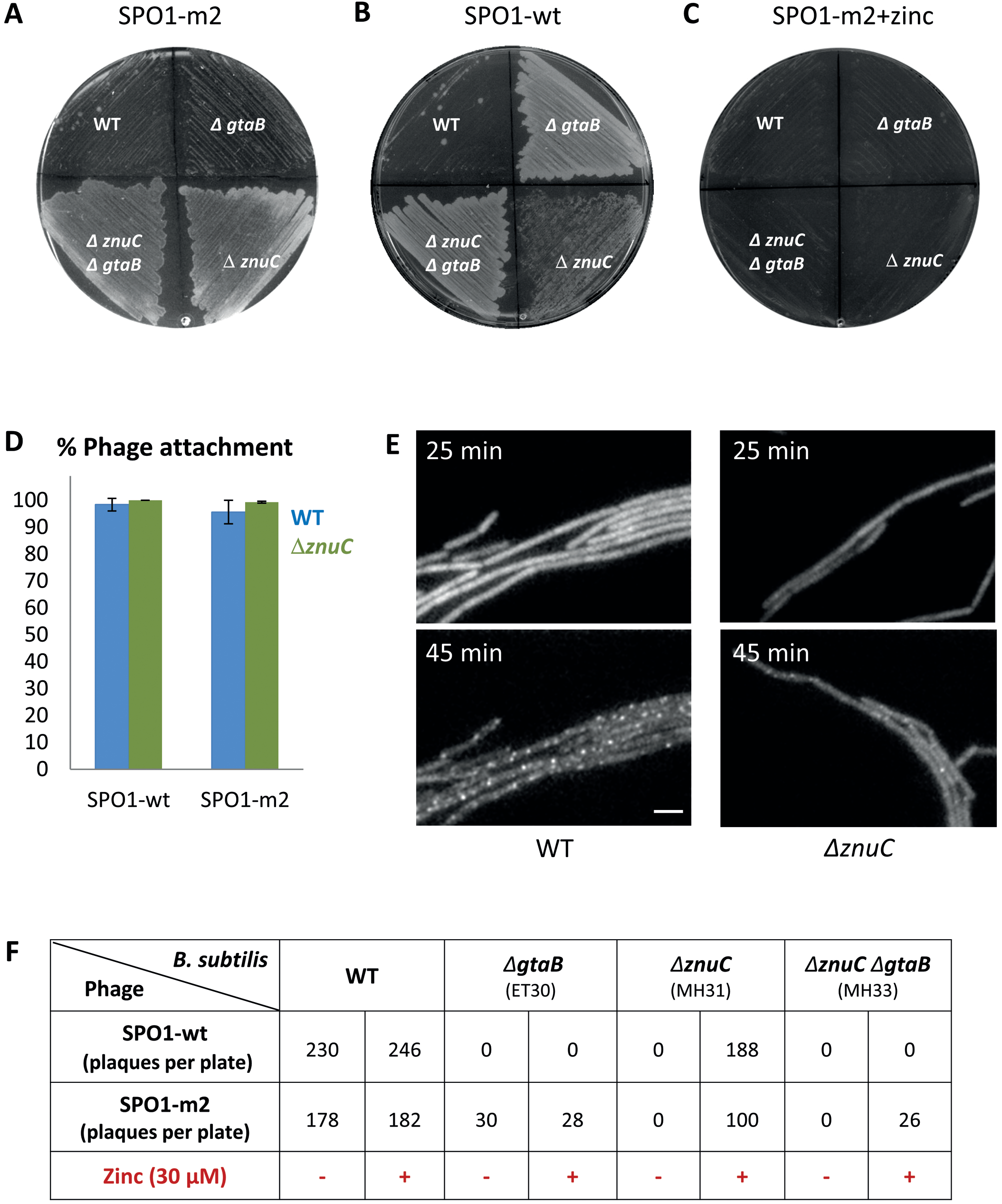
The ZnuC ABC transporter affects SPO1 infectivity. **(A-C)** Strains WT (PY79), *ΔgtaB* (ET30), *ΔznuC* (MH31) and *ΔgtaB ΔznuC* (MH33) were streaked over an agar plate containing (A) SPO1-m2, (B) SPO1-wt, and (C) SPO1-m2, supplemented with 30 μM zinc. Plates were photographed after 20 hrs of incubation at 37°C. Zinc supplementation suppressed the resistance phenotype to SPO1-m2 observed by *ΔznuC* cells. **(D)** WT (PY79) and *μznuC* (MH31) strains were infected with SPO1-wt or SPO1-m2 (MOI=0.1) and phage adsorption was monitored after 12 min by PFU/ml. Percentage phage adsorption was calculated as follows: (P_0_-P_1_)×100/P_0_, where P_0_ is the initial phage input in the lysate (PFU/ml), and P_1_ is the titer of free phages (PFU/ml) after cell infection. Shown are average values and SD of at least 2 independent experiments. **(E)** WT (ET8) or *ΔznuC* (MH35) cells harboring P_xyl_-*gfp*-*gp6.1* were grown in the presence of xylose (0.25%), infected with SPO1-wt (MOI=10) (t=0), and followed by time lapse microscopy. Shown are fluorescence images from GFP-GP6.1 at the indicated time points post infection. Scale bar, 2 μm. **(F)** SPO1-wt and SPO1-m2 phages were utilized to infect lawns of the indicated strains with or without the presence of 30 μM zinc, and infection efficiency was detected by plaque number. Shown is a representative experiment out of 3 independent repeats.

**Figure S4.**
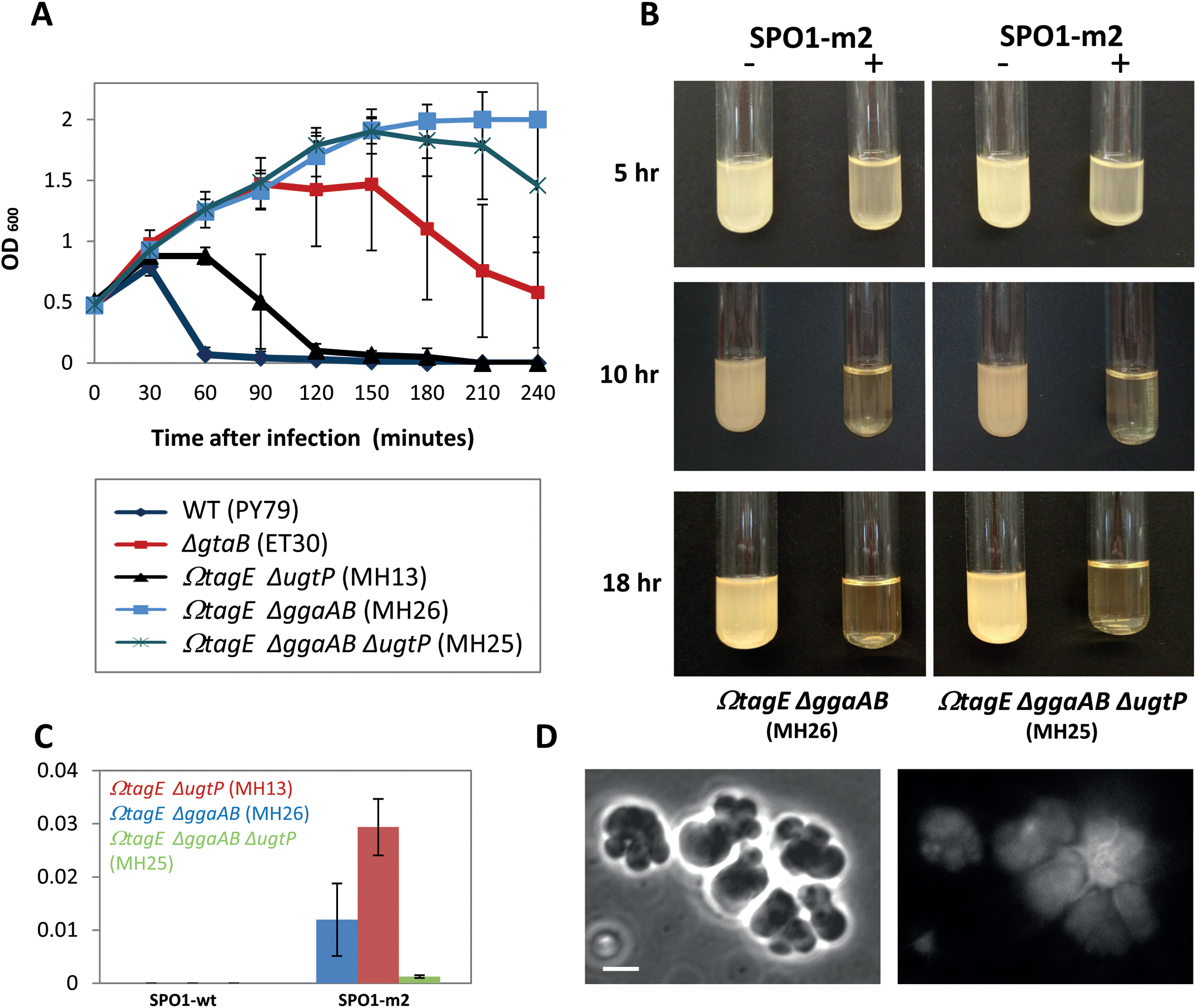
SPO1-m2 infection of various TA mutant strains. A. Lysis assay of WT (PY79) and the indicated mutant strains infected with SPO1-m2 (MOI=10) (t=0). Lysis was followed by OD_600_ at the indicated time points. Shown are average values and SD of at least 3 independent experiments. Values for ET30 and MH25 are an average of 13 and 5 independent experiments respectively, due to variability in cell lysis time.
B. MH26 (Ω*tagE* Δ*ggaAB*) (left panels) and MH25 (Ω*tagE* Δ*ggaAB* Δ*ugtP*) (right panels) strains were grown in the absence (-) or the presence (+) of SPO1-m2 (MOI=10), and cultures were photographed at the indicated time points.
C. WT (PY79) or the indicated mutant strains were infected with the indicated phages at the same PFU, and the number of plaques was monitored after 20 hrs. Shown is EOP (efficiency of plating) on the various mutants: [number of plaques on each mutant/ number of plaques on WT]. Results are average values and SD of at least 3 independent experiments.
D. Shown are phase contrast (left panel) and DAPI staining (right panel) of MH36 (Δ*tagO*) cells during logarithmic phase.

**Figure S5.**
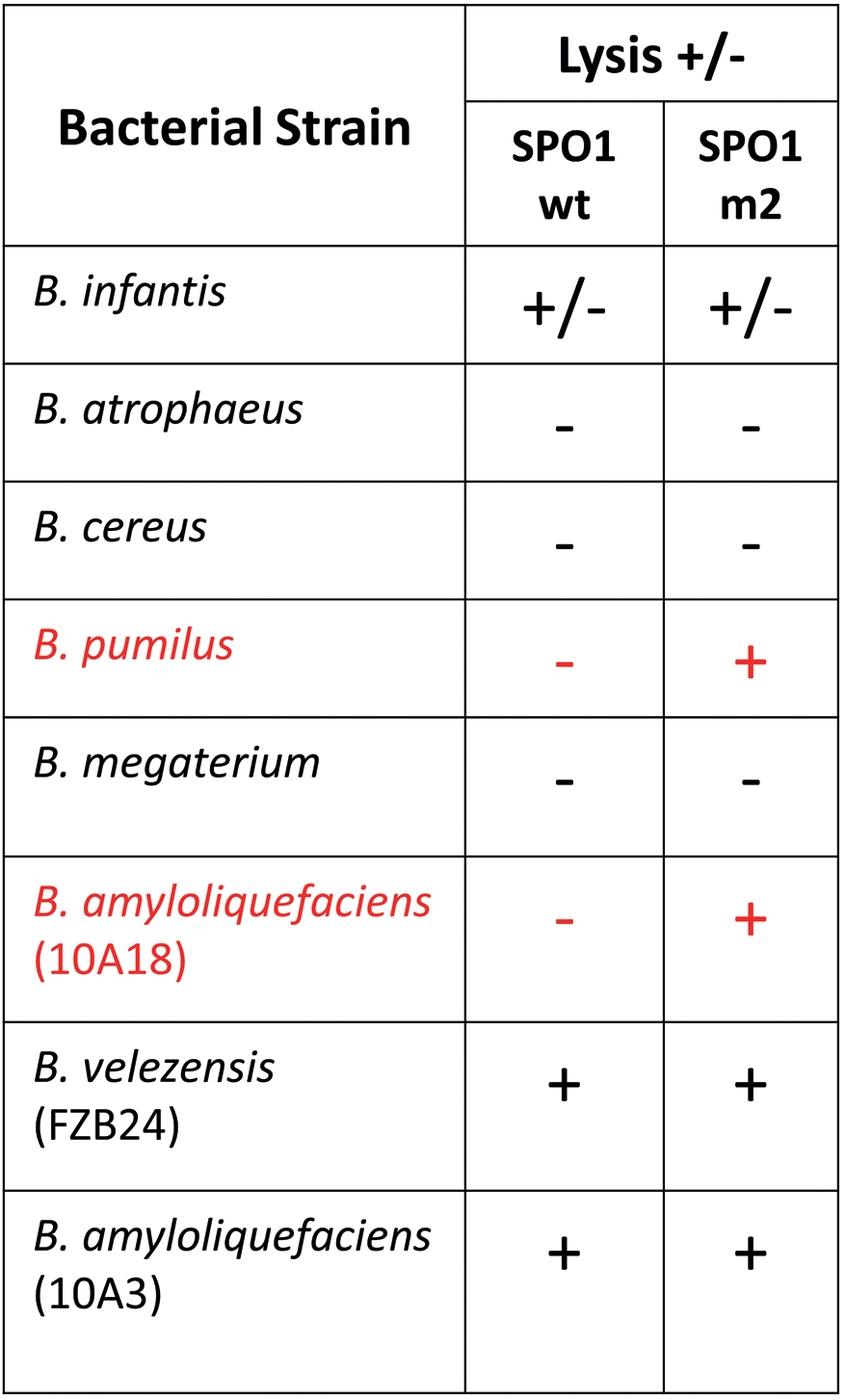
Investigating the capability of SPO1-m2 to infect various species. SPO1-wt and SPO1-m2 phages (10^8^ PFU) were spotted on lawns of the listed strains and cell lysis was observed after overnight. *B. amylo* (10A18) and *B. pumilus*, highlighted in red, could be lysed only by SPO1-m2.

**Figure S6.**
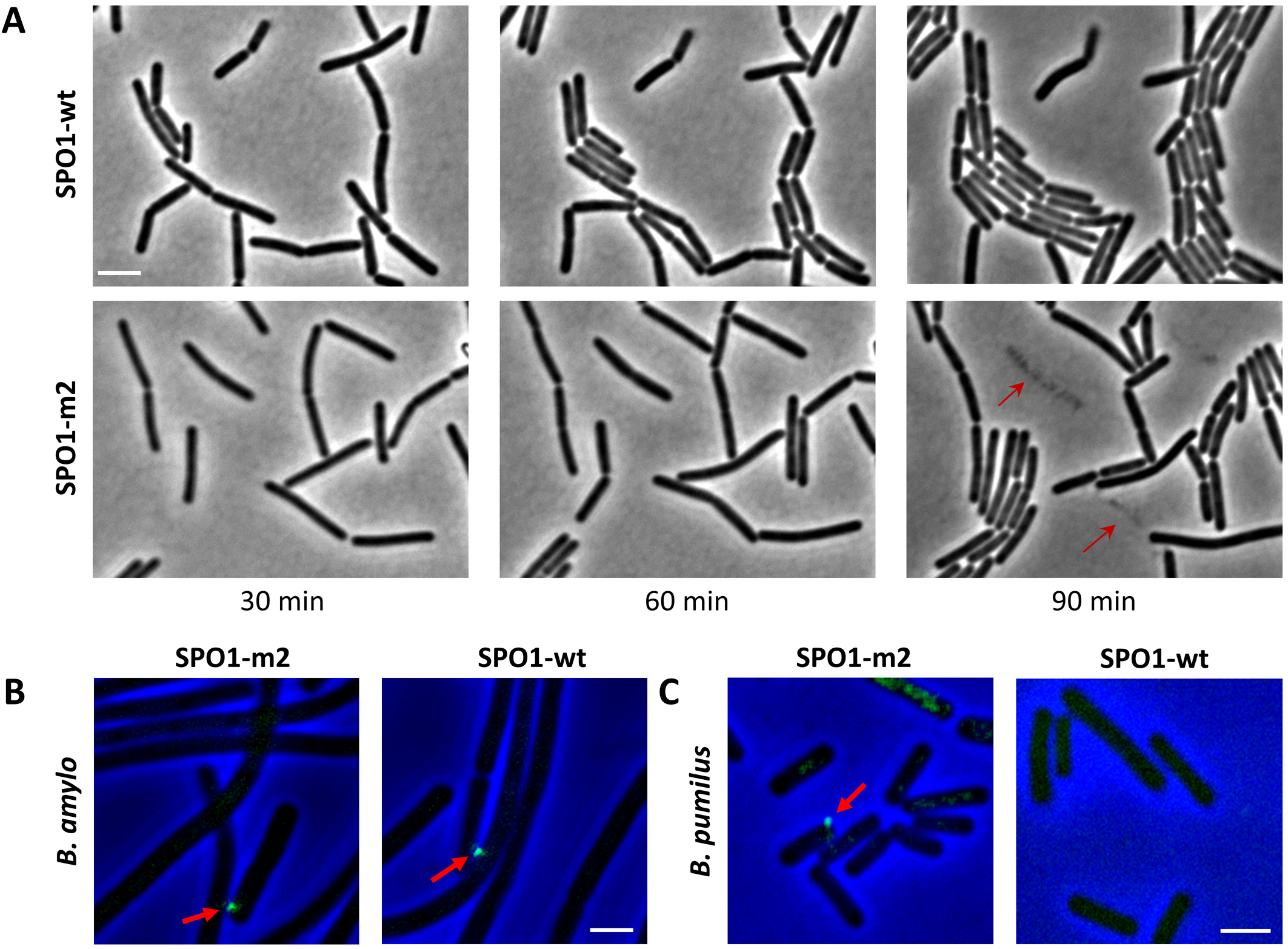
Spo1-m2 can infect non-host strains. **(A)** Infected *B. pumilus* cells were followed by time lapse microscopy. Shown are phase contrast images taken at the indicated time points post infection (t=0) with SPO1-wt (upper panels) or SPO1-m2 (lower panels). Arrows highlight lysing bacterial cells. **(B-C)** SPO1-m2 (left panels) and SPO1-wt (right panels) were labeled with DAPI and phage attachment to *B. amylo* (10A18) (B) or *B. pumilus* (C) was detected by fluorescence microscopy. Shown are overly images of phase contrast (blue) and DAPI fluorescence (green). Arrows indicate phage binding to the cell surface. Scale bars, 2 μm.

